# Benchmarking confidence estimation and rescoring for cyclic peptide–protein complex predictions

**DOI:** 10.64898/2026.08.20.746104

**Authors:** Zhe Li, Ye Yuan, Kaiqiang Hu, Pengwei Pan, Fang He

## Abstract

Cyclic peptides are a rapidly expanding class of therapeutics, but the reliability of deep-learning structure prediction for cyclic peptide–protein complexes has not been systematically evaluated. We assembled a curated benchmark of 111 non-redundant complexes spanning five cyclization chemistries and assessed two co-folding models, Boltz and Protenix, each generating 100 poses per target (22,200 total). Stratifying all poses by complex attributes, we found that disulfide-cyclized peptides and small protein targets (200 or fewer target residues) were predicted significantly worse by both tools, with target size the largest and most consistent effect; overall accuracy nevertheless remained high (median top-pose DockQ of about 0.89, 96–98% of targets Acceptable or better), indicating that pose generation is rarely the bottleneck. Conversely, native model ranking scores correlated only moderately with pose quality (Spearman rank correlations of 0.53–0.66): approximately 12% of poses showed high model ranking score/confidence despite poor pose DockQ quality, and the highest-quality pose was not ranked first for nearly every target. We therefore augmented the native score with externally computed interface descriptors normalized by chain length, principally the per-residue density of inter-chain hydrogen bonds, in a gradient-boosted rescoring model evaluated under target-grouped cross-validation that prevents leakage, improving out-of-fold ROC-AUC for both tools, significantly so for Protenix. Together, these findings identify pose ranking, rather than pose generation, as the major limitation of current cyclic peptide–protein complex prediction and demonstrate that complementary structural features can improve confidence-based pose selection.

## Introduction

Cyclic peptides have emerged as a premier modality in modern drug discovery, combining the high affinity and target specificity of biologics with structural rigidity and metabolic stability reminiscent of small molecules (Ji et al., 2024). Their constrained macrocyclic framework reduces the entropic cost of binding, enhances proteolytic resistance, and enables engagement of extended or shallow protein surfaces that are inaccessible to conventional small-molecule inhibitors (Ji et al., 2024; Zhang and Chen, 2022). Between 2001 and 2021 alone, 18 cyclic peptide drugs received regulatory approval worldwide, treating indications from antimicrobial therapy and immunosuppression to oncology, and the development pipeline continues to expand rapidly (Zhang and Chen, 2022). Rational optimization of these molecules (whether to improve binding affinity, selectivity, or pharmacokinetic properties) depends critically on accurate three-dimensional structures of the cyclic peptide–protein complex, which reveal the intermolecular contacts, hydrogen-bond networks, and conformational constraints that govern recognition.

Building on the AlphaFold line of structure-prediction models, fully open-source co-folding engines such as Boltz (Wohlwend et al., 2024; Passaro et al., 2025) and Protenix (Protenix Team et al., 2026) now predict arbitrary biomolecular assemblies at all-atom resolution, including covalent modifications. Importantly for cyclic peptides, these tools accept user-specified covalent constraints (head-to-tail backbone cyclization and disulfide bridges), and recent studies have demonstrated that computational and deep-learning models can fold cyclic peptides with varying degrees of success (Rettie et al., 2025; Xie et al., 2025; Liu et al., 2020).

Structure prediction models are best thought of as conformation samplers. Indeed, selecting the single most reliable pose from a sampled ensemble is arguably the most critical task in downstream applications, including structure-based drug design (SBDD). Both Boltz and Protenix, following the AlphaFold paradigm, rank predicted structures using internal confidence metrics derived from predicted aligned error and inter-chain predicted TM (ipTM) scores (Abramson et al., 2024; Zhai et al., 2025). These confidence estimates have become the de facto selection criterion in virtually all structure-prediction workflows. However, their reliability for cyclic peptide–protein complexes (where the cyclic constraint and frequently shallow binding interfaces may confound the coevolutionary and distance-prediction signals on which these scores depend) has not been systematically evaluated. Recent benchmarking of AlphaFold 3 for linear protein–peptide complexes has revealed substantial overconfidence and pose-selection failures (Genz et al., 2025; Peng et al., 2025; Zhai et al., 2025), but it remains unknown whether and to what extent these limitations extend to cyclic-peptide systems.

More broadly, existing research lacks a comprehensive benchmark specifically addressing confidence estimation for cyclic peptide–protein complex prediction. Prior benchmarking efforts have evaluated traditional docking and rescoring methods for cyclic peptides (Zhao et al., 2024, 2025) or focused on linear peptide folding and complex prediction (Zhai et al., 2025), and none have systematically examined how cyclization chemistry (head-to-tail backbone cyclization, disulfide bridging, or both) influences confidence-score reliability across a diverse, non-redundant target set. Consequently, it remains unclear under what circumstances current confidence metrics fail for cyclic peptides, which structural or chemical features of the complex are associated with overconfident or underconfident predictions, and whether externally computed interface descriptors can complement native confidence to improve pose ranking.

To address these gaps, we assembled a curated benchmark of 111 experimentally determined cyclic peptide– protein complex structures spanning five cyclization chemistries (head-to-tail backbone cyclization; head-to-tail backbone cyclization combined with one or more intra-peptide disulfide bridges; and peptides cyclized solely by one to three disulfide bridges), and covering a broad range of binder and target sizes. Using this benchmark, we systematically evaluated the confidence-estimation performance of Boltz and Protenix, each generating 100 conformers per target (22,200 poses in total), with per-pose quality scored by DockQ v2 in CAPRI-peptide mode (Basu and Wallner, 2016; Mirabello and Wallner, 2024). We characterized the common failure modes of native confidence scores, including overconfident wrong poses and missed high-quality poses within sampled ensembles, and analyzed how these failures distribute across cyclization types, binder lengths, and target sizes. Finally, we developed a machine-learning-based confidence-refinement strategy that augments each model’s native self-confidence with externally computed interface descriptors normalized by chain length (including hydrogen-bond density, interface burial, and electrostatic features), demonstrating consistent improvements in cross-target pose ranking for both prediction tools. Together, our study shows that the principal limitation of current cyclic peptide– protein complex prediction lies not in generating near-native structures but in ranking them, and that simple, physically interpretable interface descriptors, principally per-residue inter-chain hydrogen-bond density, can materially improve confidence-based pose selection.

## Results

### A curated benchmark and the target attributes that govern prediction accuracy

To assemble a chemistry-aware benchmark for cyclic peptide–protein complex prediction, we integrated previously published cyclic peptide–protein complex data (Zhao et al., 2024, 2025) with publicly available Protein Data Bank structures. Evaluating structure-prediction methods on cyclic peptides is hampered by the absence of a clean, non-redundant benchmark: public sets routinely mix backbone- and disulfide-cyclized chemistries, retain redundant crystallographic copies of the peptide or receptor, and span several orders of magnitude in binder and target length. To attribute prediction accuracy to chemically meaningful features of the complex rather than to dataset artifacts, we assembled a curated benchmark of 111 non-redundant cyclic peptide–protein complexes, composed exclusively of natural amino acids (entries containing non-canonical residues were excluded), standardized to a single primary conformer and a single binder–receptor complex per target (Figure 1). The set spans the full spectrum of cyclic-peptide chemistries, namely head-to-tail backbone cyclization (BB, n = 27), backbone cyclization combined with one or more intra-peptide disulfide bridges (BB+SS, n = 10), and disulfide-only peptides with one to three bridges (SS*1, n = 61; SS*2, n = 12; SS*3, n = 1), and a broad range of binder (6–34 residues) and target (67–902 residues) sizes (target length defined as the total number of residues in the receptor chains contacted by the binder; see Methods). This composition provides the stratification axes needed to dissect performance.

**Figure 1.**
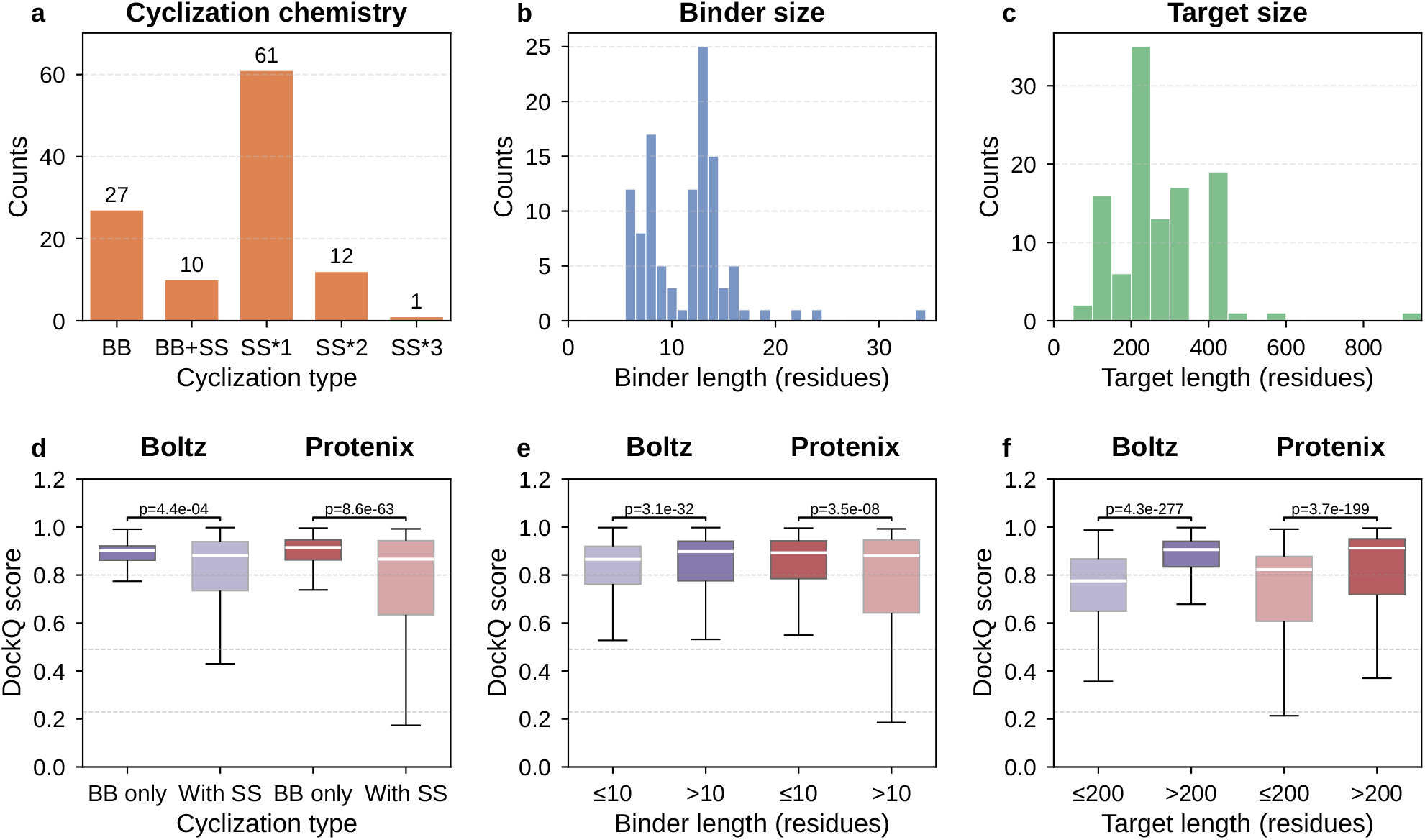
Dataset composition and prediction-quality stratification for the curated cyclic peptide–protein benchmark. **(a–c)** Composition of the curated dataset (n = 111 complexes). **(a)** Cyclization chemistry, classified as backbone-cyclized (BB, head-to-tail N(1)–C(last) amide bond), backbone plus one or more disulfides (BB+SS), or disulfide-only peptides with one, two, or three disulfide bridges (SS*1, SS*2, SS*3); counts are annotated above each bar. **(b)** Distribution of cyclic-peptide (binder) length, ranging from 6 to 34 residues. **(c)** Distribution of target length, defined as the total number of residues in the receptor chains contacted by the binder (see Methods), ranging from 67 to 902 residues. **(d–f)** Distributions of pose-level CAPRI-peptide DockQ scores for Boltz (purple) and Protenix (red) across all 111 complexes (100 poses each; 22,200 poses total), with no per-target aggregation; within-tool class differences are tested with a two-sided Mann–Whitney U test (p-value above each bracket). Dashed lines mark the DockQ High (0.80), Medium (0.49), and Acceptable (0.23) tiers. **(d)** Cyclization class: backbone-only (BB) vs disulfide-cyclized (SS). **(e)** Binder length: 10 or fewer vs more than 10 residues. **(f)** Target length: 200 or fewer vs more than 200 target residues (definition as in **c**).

We then evaluated two state-of-the-art co-folding models, Boltz (Passaro et al., 2025) and Protenix (Protenix Team et al., 2026), each generating 100 conformers per target (10 seeds × 10 samples; 22,200 poses in total), and scored every pose against its native structure with DockQ v2 in CAPRI-peptide mode (Basu and Wallner, 2016; Mirabello and Wallner, 2024). On the primary (top-confidence) pose both tools already perform well, reaching median CAPRI-peptide DockQ of 0.888 (Boltz) and 0.896 (Protenix) across the 111 targets, with 96–98% of targets scoring Acceptable or better. To understand where the remaining errors concentrate, we stratified all 22,200 poses by three binary attributes of the complex (cyclization chemistry, binder length, and receptor size) and compared the pose-level DockQ distributions within each tool, using Mann–Whitney U tests (Figure 1).

Stratifying by cyclization chemistry revealed a consistent penalty for disulfide-cyclized peptides (cyclized by disulfide bridges alone or in addition to backbone cyclization): collapsing the five classes into backbone-only (BB) versus disulfide-cyclized targets, the latter showed lower median DockQ for both Boltz (0.90 vs 0.88, p = 4.4×10^−4^) and Protenix (0.91 vs 0.87, p = 8.6×10^−63^).

This is consistent with reports that disulfide-rich peptides are refractory to AlphaFold-style folding: satisfying several cystine cross-links simultaneously, on top of the backbone-cyclization constraint, imposes geometry that co-folding models, trained predominantly on globular protein complexes, resolve poorly without explicit connectivity restraints (Xie et al., 2025; Liu et al., 2020; Gerlach and Nicoludis, 2024; Rettie et al., 2025). Our result extends this limitation to the bound complex, showing that the penalty carries over into interface placement. The binder-length effect is tool-dependent. Longer binders (more than 10 residues) were modeled slightly better by Boltz (median 0.90 vs 0.87 for 10 or fewer, p = 3.1×10^−32^), whereas Protenix favored shorter binders (0.89 vs 0.88, p = 3.5×10^−8^); both contrasts were significant but the effects were small and opposite in sign. This tool-specific behavior suggests that binder-length sensitivity reflects differences in how each model encodes the cyclic constraint and samples the peptide backbone, rather than an intrinsic difficulty of the task. Larger targets are predicted substantially better by both tools. With the split at 200 target residues, small targets were markedly harder (Boltz 0.78 vs 0.91, p = 4.3×10^−277^; Protenix 0.82 vs 0.91, p = 3.7×10^−199^); this was the largest and most consistent effect in the figure. We attribute this to two cooperating factors. Larger receptors generally present more extensive, geometrically defined binding surfaces that give the binder fewer competing registers and yield a stronger inter-chain confidence signal (ipTM), whereas small targets disproportionately present shallow, flexible, or multi-chain interfaces that are intrinsically harder to dock. In addition, larger single-chain receptors carry deeper, more informative multiple-sequence alignments (MSAs), on which the coevolution-driven confidence of both models depends (Passaro et al., 2025; Protenix Team et al., 2026).

### Native confidence scores correlate with quality but fail to surface the best pose

These accuracy trends are based on DockQ, which is unavailable at prediction time. In practice, users must select a pose using each model’s own confidence estimate: Boltz’s composite confidence score and Protenix’s ranking score (Abramson et al., 2024; Zhai et al., 2025). To ask how reliably these self-scores rank pose quality, we plotted pose-level CAPRI-peptide DockQ against the native self-confidence for every sampled pose, both across the full pool and for the single top-confidence pose per target (Figure 2).

**Figure 2.**
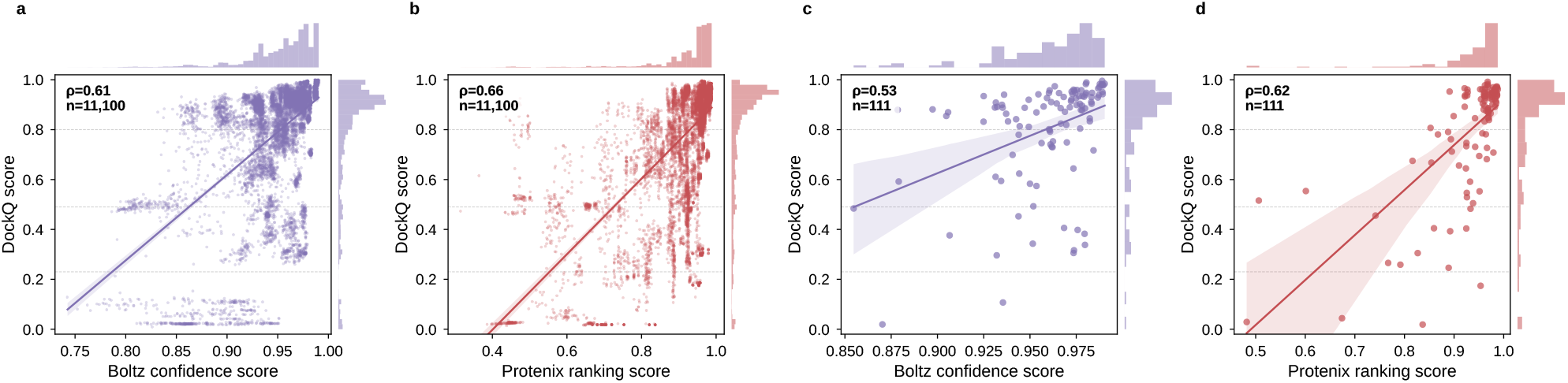
Tool self-confidence versus prediction quality. Scatter of CAPRI-peptide DockQ against each tool’s native self-confidence score (Boltz confidence score, purple; Protenix ranking score, red), with a linear regression fit (95% CI) and top/right marginal histograms. Dashed lines mark the DockQ High (0.80), Medium (0.49), and Acceptable (0.23) tiers. **(a, b)** All 11,100 poses per tool (10 seeds × 10 samples × 111 complexes); Spearman ρ = 0.61 (Boltz) and 0.66 (Protenix). **(c, d)** The single highest-confidence pose per target (n = 111), i.e. each tool’s own top-confidence pick; Spearman ρ = 0.53 (Boltz) and 0.62 (Protenix). Scores are shown on each tool’s native [0, 1] scale.

Overall, self-confidence and DockQ are positively correlated: across all 11,100 poses per tool the Spearman correlation is ρ = 0.61 for Boltz and ρ = 0.66 for Protenix, consistent with the general finding that ipTM-based scores carry useful, if imperfect, information about interface quality (Genz et al., 2025; Peng et al., 2025). The relationship is far from one-to-one, however. A substantial fraction of poses are overconfident, combining a high self-score (0.5 or higher) with a sub-Medium DockQ (below 0.49): about 12% of poses for both tools (11.7% Boltz; 12.9% Protenix). The cloud of points in Figure 2a,b therefore contains a clear lower-right population of confidently wrong predictions, and concentrating on each model’s own top pick per target (Figure 2c,d) only weakens the correlation (ρ = 0.53 and 0.62, n = 111) rather than sharpening it; the self-score is a noisy ranker precisely where selection matters most. This selection failure is hidden by the favorable aggregate statistics. Although the top-confidence pose is Acceptable-or-better for 96–98% of targets, it is not the best available pose in almost every case: comparing each tool’s confidence pick against the best pose within its own 100-pose pool, 108 of 111 (Boltz) and 110 of 111 (Protenix) targets retain a positive quality gap (one-sided exact sign tests, p = 8.8×10^−29^ and 4.3×10^−32^, respectively), and the gap reaches 0.69–0.91 DockQ at its extreme. In other words, a pose of higher quality than the model’s own pick is present for 97–99% of targets, but native confidence does not reliably identify it, a limitation repeatedly noted for ipTM-style scoring of multimeric predictions (Genz et al., 2025; Peng et al., 2025).

To further illustrate this gap, we selected four representative targets and visualized their full pose-confidence ladders: all 100 sampled poses at (self-score, DockQ), with the model’s confidence pick (open ring), the best pose in the pool (star), and the selection arrow between them (Figure 3). Two targets (5H5R, 3AVC) are solved reliably by both tools, for which the confidence pick already coincides with a near-best pose. The other two were chosen because their confidence behavior flips across tools, isolating the selection failure from any intrinsic difficulty of the target: on 5VB9, Boltz is overconfident (its top-confidence pick scores DockQ 0.11 while the best-scoring pose reaches 0.83; Δ = 0.72), whereas Protenix appropriately assigns low confidence and selects a near-best pose; on 4GLY the pattern reverses, with Protenix overconfident (pick 0.17, best 0.68; Δ = 0.50) and Boltz reliable. In both flip cases the model assigns its highest confidence to a wrong register of the same cyclic peptide, while a correct pose is present among the sampled conformers. Figure 3i–l visualizes these top-confidence picks in 3D against their native complexes, making the register agreement (5H5R, 3AVC) and the wrong-register displacement in the two flip cases directly visible.

**Figure 3.**
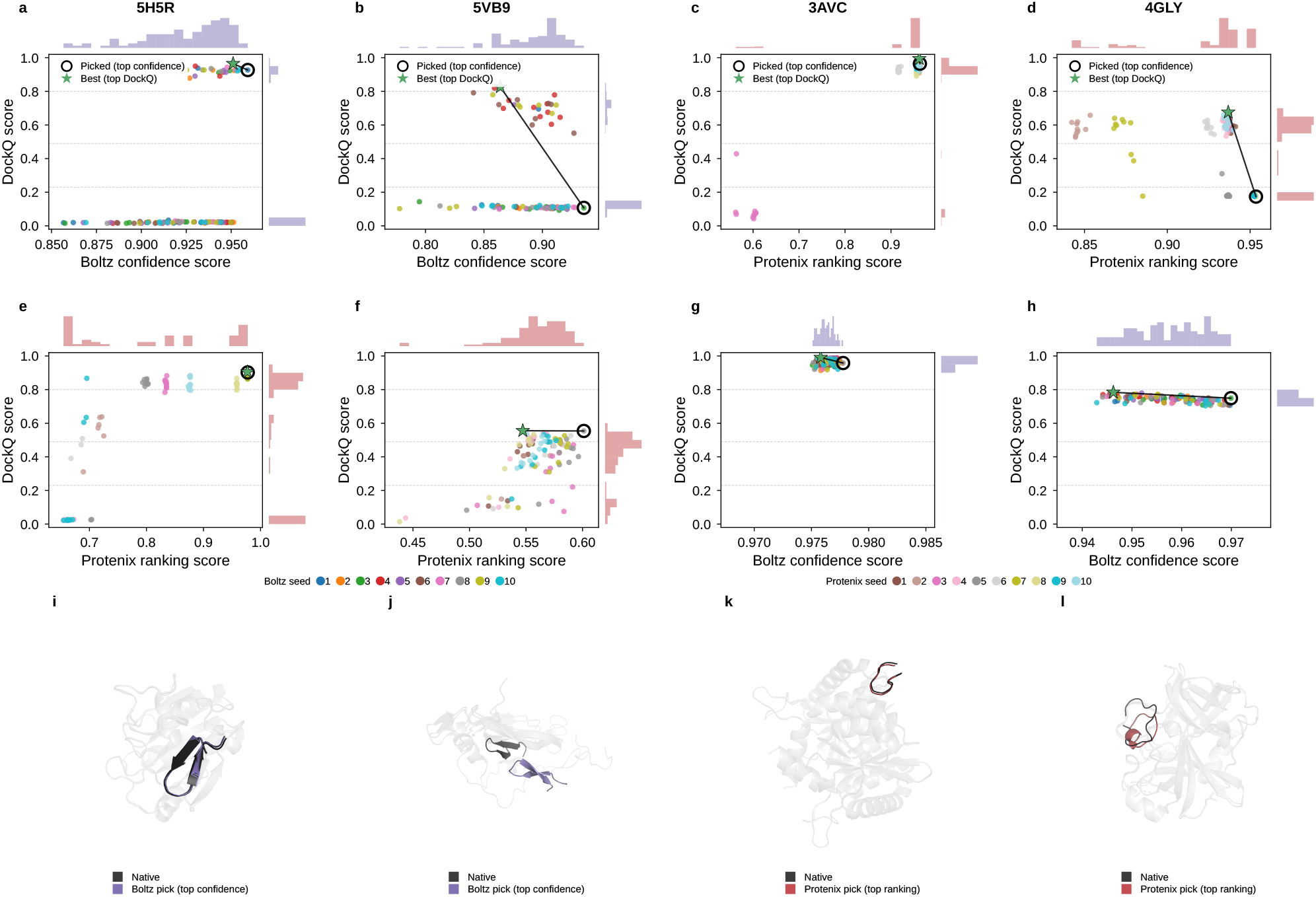
Pose–confidence ladders for representative targets. Each panel shows all 100 sampled poses (10 seeds × 10 samples; points colored by seed) for one tool–target combination, plotting CAPRI-peptide DockQ against the tool’s native self-confidence score (Boltz confidence score or Protenix ranking score). The open ring marks the tool’s own top-confidence pick, the green star marks the highest-DockQ pose in the pool, and the black arrow (drawn where they differ) exposes the selection gap. Dashed lines mark the DockQ High (0.80), Medium (0.49), and Acceptable (0.23) tiers; top/right marginal histograms show the score distributions. The 2×4 grid pairs the same target across tools by column: 5H5R **(a, e)**, 5VB9 **(b, f)**, 3AVC **(c, g)**, 4GLY **(d, h)**. 5H5R and 3AVC succeed for both tools; 5VB9 and 4GLY flip, with Boltz overconfident on 5VB9 and Protenix overconfident on 4GLY (large selection gaps in **b** and **d**), while the respective other tool selects a near-best pose. **(i–l)** Structural view of each column’s top-confidence pick: the predicted cyclic peptide (colored) superposed on the native complex (gray translucent targets; near-black native binder) by receptor Cα alignment. On 5H5R and 3AVC **(i, k)** the predicted binder overlays the native peptide at the same binding register; on 5VB9 and 4GLY **(j, l)** the picked pose occupies a wrong register of the same peptide, visible as the displaced colored ring.

### External interface descriptors complement native confidence and improve pose ranking

The preceding analysis showed that each model’s native confidence is only loosely coupled to pose quality. Nor do the models’ other internal metrics offer an alternative: the redundancy structure of the full feature pool (the model-internal signals together with the chain-length-normalized interface descriptors), visualized as pooled per-pose Spearman correlation heatmaps (Figure 4), shows that the model-internal signals (pTM, ipTM, pLDDT, and the PDE-type distances) form a single, tightly inter-correlated block (a “confidence axis”, with correlations to the composite self-score of |ρ| = 0.72–1.00; e.g., Protenix ipTM vs ranking score, ρ = 0.996), so none of them carries information substantially different from the native self-score. We therefore asked whether externally computed interface descriptors carry information that this confidence axis misses, and whether folding that information into a model can rescue pose selection. In contrast to the internal block, the external interface descriptors are largely decoupled from the confidence axis and from one another, providing a pool of candidate signals that native confidence does not capture. Decoupling alone, however, does not establish usefulness: Protenix’s target net charge is essentially orthogonal to the ranking score (ρ of roughly 0.04), whereas the per-residue hydrogen-bond densities and Boltz’s SASA per target residue are moderately correlated with the confidence axis (|ρ| = 0.32–0.51); their value lies in incremental predictive signal, which we test next.

**Figure 4.**
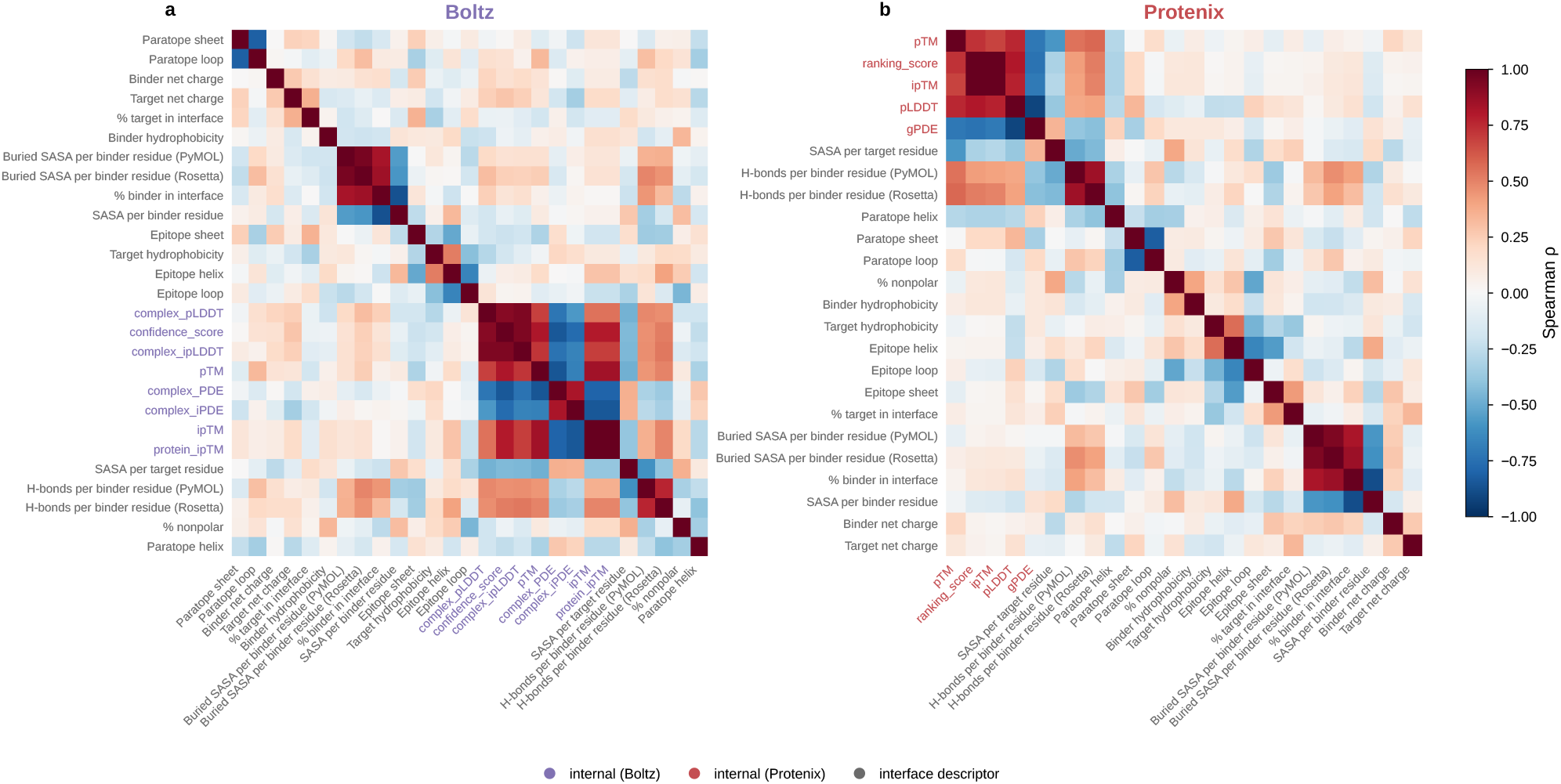
Correlation of model-internal and interface features. Per-tool Spearman rank-correlation heatmaps over all 11,100 poses (111 targets × 10 seeds × 10 samples) between model-internal confidence signals and a chain-length-normalized set of interface descriptors. **Internal signals** (model-emitted) are 8 for Boltz (confidence score, pTM, ipTM, complex pLDDT, protein ipTM, complex ipLDDT, PDE, iPDE) and 5 for Protenix (ranking score, pTM, ipTM, pLDDT, gPDE). **Interface descriptors** (computed by PyMOL/Rosetta on the predicted structure, or from sequence) comprise 12 already-comparable features (“% nonpolar” (% of interface residues that are non-polar); “Paratope helix/sheet/loop” and “Epitope helix/sheet/loop” (% of binder-side and target-side interface residues in each secondary-structure class); “% binder in interface”; “Binder/Target hydrophobicity” (mean sequence hydrophobicity); “Binder/Target net charge”) and 7 chain-length-normalized densities: “SASA per binder residue” and “SASA per target residue” (total solvent-accessible surface area per chain residue); “Buried SASA per binder residue (PyMOL)” and “(Rosetta)” (interface buried SASA per binder residue, from each tool); “% target in interface”; and “H-bonds per binder residue (PyMOL)” and “(Rosetta)” (inter-chain hydrogen bonds per binder residue). **(a)** Boltz. **(b)** Protenix. Cells are pooled per-pose Spearman ρ on a diverging scale (−1 to +1); features are reordered by average-linkage clustering on distance 1 - |ρ| so that strongly correlated (positively or negatively) features cluster adjacent. Tick labels are color-coded: internal signals in the tool color (Boltz purple, Protenix red) and interface descriptors in gray.

To quantify their predictive value, we trained a gradient-boosted classifier to label each pose as Good (CAPRI-peptide DockQ above 0.80) or Bad, evaluated under target-grouped cross-validation (CV) that prevents leakage, with the native self-score force-seeded into every model and the remaining descriptors selected by Sequential Forward Floating Selection (SFFS (Pudil et al., 1994); Figure 5; Methods). The univariate separation underlying this classification is shown in Figure S3: across all 22,200 poses, z-scored per-feature boxplots confirm that model-internal confidence signals and, among interface descriptors, per-residue inter-chain hydrogen-bond density separate Good from Bad most strongly for both tools (Good/Bad counts of 7,904/3,196 for Boltz and 7,261/3,839 for Protenix). The consensus model was assessed at three levels: the inner-CV ROC-AUC trajectory traced during SFFS feature addition (Figure 5a,b), the pooled out-of-fold ROC contrasting the self-score baseline with the feature-augmented model (Figure 5c,d), and the TreeSHAP contributions of the selected descriptors (Figure 5e,f).

**Figure 5.**
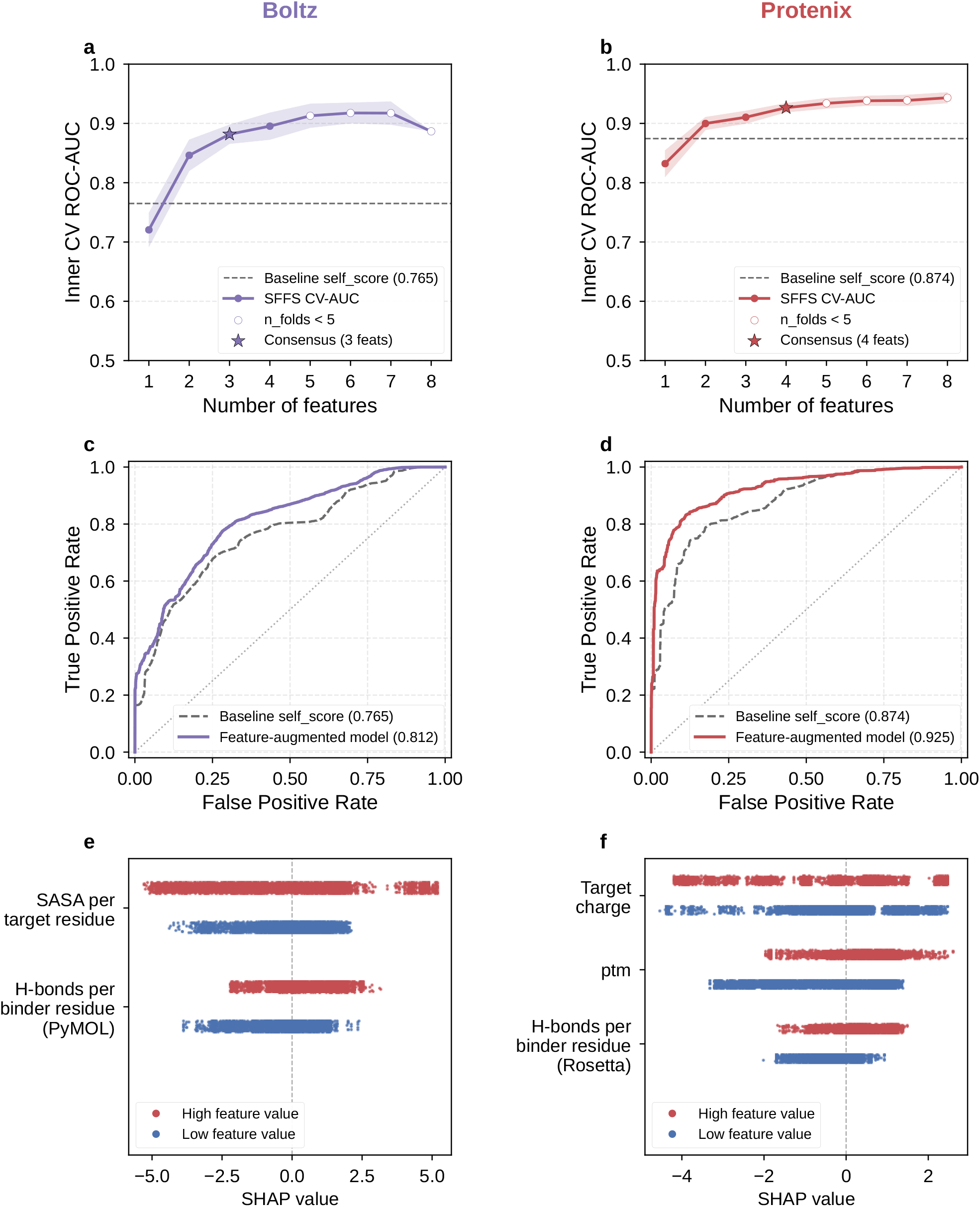
Learning to predict per-pose quality. A LightGBM binary classifier is trained to label each predicted pose as Good (CAPRI-peptide DockQ above 0.80) or Bad, using the chain-length-normalized feature set of Figure 4 with each tool’s self-confidence score force-seeded. Evaluation is leakage-free by target (StratifiedGroupKFold(5) grouped by pdb_id over 11,100 poses: 111 targets × 10 seeds × 10 samples), with Sequential Forward Floating Selection (SFFS) in an inner 5-fold loop. Columns are Boltz (purple) and Protenix (red). **(a, b)** SFFS inner-CV ROC-AUC versus number of features; the dashed line marks the self_score-only baseline and the star the consensus feature count. **(c, d)** Out-of-fold ROC curves of the baseline (self_score alone) versus the feature-augmented model; AUC rises from 0.765 to 0.812 for Boltz and from 0.874 to 0.925 for Protenix. **(e, f)** SHAP split-strip plots of the SFFS-selected features (Boltz: “SASA per target residue” and “H-bonds per binder residue (PyMOL)”; Protenix: “Target charge”, “ptm” and “H-bonds per binder residue (Rosetta)”): for each feature, poses with high and low feature values (red and blue, respectively) are shown as two solid-color strips at different vertical offsets within the feature row, so the two distributions never overlap; strip position along the horizontal axis shows each pose’s SHAP contribution to the Good-pose prediction.

Augmenting the self-score with a small number of selected descriptors improved cross-target ranking for both tools, though with different statistical confidence. Out-of-fold ROC-AUC rose from 0.765 to 0.812 for Boltz (ΔAUC = 0.046, 95% CI −0.036 to 0.128, p = 0.28) and from 0.874 to 0.925 for Protenix (ΔAUC = 0.051, 95% CI 0.014 to 0.093, p < 0.001; 1,000-resample target-level block bootstrap in both cases): the Protenix gain is statistically significant, whereas the Boltz gain, similar in magnitude, is not individually significant, consistent with its wider confidence interval. The improvement is modest in absolute terms (self-confidence is genuinely hard to beat for cross-target Good/Bad prediction), but its direction is consistent across tools and confirms that the external descriptors carry real signal beyond what ipTM-based scores already provide.

The feature-contribution panels point to a shared, physically interpretable signal (Figure 5e,f). The single most useful complementary descriptor for both tools is the per-residue inter-chain hydrogen-bond density (H-bonds per binder residue, from PyMOL for Boltz and Rosetta for Protenix). Interface hydrogen bonds are a direct readout of satisfied polar contacts and shape/electrostatic complementarity at the binding surface, and are established determinants of interface quality and complex stability that coarse-grained coevolutionary confidence scores tend to under-weight (Kuroda and Gray, 2016; Pace et al., 2014). The length-normalized, per-residue form of this descriptor makes it comparable across binders of different lengths, which is why it survives SFFS where raw H-bond counts do not. The two variants of this descriptor encode different definitions of the same physical quantity (PyMOL’s geometric distance–angle criteria versus Rosetta’s knowledge-based H-bond energy term) and are strongly but not identically correlated (ρ = 0.77 for Boltz poses, 0.85 for Protenix poses), with the Rosetta variant separating Good from Bad slightly better in univariate analysis (ρ = 0.33–0.36 vs 0.24–0.27). SFFS retains exactly one variant per tool (PyMOL for Boltz, Rosetta for Protenix) because once either enters the model the other is redundant; the tool-specific choice thus reflects collinearity-driven selection among near-equivalent features rather than a principled preference for one H-bond definition. Likewise, the PyMOL and Rosetta buried-SASA densities are nearly identical (ρ = 0.94–0.95) and neither was selected. The remaining selected features are tool-specific: for Boltz, SASA per target residue contributes a measure of relative interface burial (another known correlate of complex stability (Chakravarty et al., 2013)) that is anti-correlated with the confidence axis and thus supplies an independent failure signal; for Protenix, target net charge adds electrostatic-complementarity information that is nearly orthogonal to the ranking score, while pTM contributes intra-chain folding confidence that complements the inter-chain ranking score.

## Discussion

This study presents a systematic, chemistry-stratified evaluation of confidence estimation for cyclic peptide– protein complex prediction. Our central finding is that the native confidence scores of Boltz and Protenix, while positively correlated with pose quality, are too noisy to serve as reliable sole ranking criteria for cyclic peptide–protein poses. The per-target correlations between self-confidence and DockQ were only moderate (ρ = 0.53–0.62), approximately 12% of all poses were overconfident, and the best available pose was missed for nearly every target. These patterns are consistent with recent reports of ipTM overconfidence for AlphaFold 3 predictions of linear peptide–protein complexes (Genz et al., 2025; Peng et al., 2025), and extend them to the cyclic-peptide regime. Importantly, the failure mode is not an inability to generate correct structures (both tools achieved median top-pose DockQ above 0.88), but an inability to identify the correct structure post hoc. The confidence metrics shared across AlphaFold-family architectures are designed to capture global inter-chain topology (ipTM) rather than residue-level binding register (Evans et al., 2021; Abramson et al., 2024), and this design objective creates a calibration gap that becomes acute for cyclic peptides: their interfaces are relatively small, so register shifts and side-chain packing errors that go undetected by a global similarity score can still dominate the local binding geometry.

The most consistent structural correlate of prediction difficulty was cyclization chemistry: disulfide-cyclized peptides were predicted significantly worse than backbone-only peptides by both tools. This effect is robust (it appears in two independent models across 84 disulfide-cyclized versus 27 backbone-only targets) and aligns with prior reports that disulfide-rich cyclic peptides are refractory to AlphaFold-style folding without explicit connectivity restraints (Xie et al., 2025; Liu et al., 2020; Gerlach and Nicoludis, 2024; Rettie et al., 2025). In our view, this penalty most plausibly reflects two non-exclusive and presently indistinguishable mechanisms. The first is a conformational-constraint mechanism: simultaneously satisfying multiple cystine cross-links and (in BB+SS targets) a head-to-tail amide bond defines a highly restricted region of backbone conformational space that iterative diffusion modules, which enforce such connectivity only through soft potentials rather than dedicated bond-constraint solvers, may fail to sample consistently. The second concerns training-data representation: deep-learning structure predictors are trained predominantly on linear, globular protein chains from the PDB, and cyclic peptides, particularly those with multiple disulfide bridges, are rare in these training corpora, limiting the models’ learned conformational priors. Recent work showing that AlphaFold 3 accuracy degrades for non-canonical cyclic peptides and that its confidence metrics are poorly calibrated for this chemistry (Zhang et al., 2025) lends circumstantial support to this representation-based view. Disentangling these mechanisms (for example, through controlled retraining with augmented cyclic-peptide data or explicit disulfide parameterization) is an important direction for future model development but is beyond the scope of this benchmarking study.

A striking implication of our pose-level analysis is that the confidence-ranking failure is predominantly a scoring problem rather than a sampling problem. For the great majority of targets in both tools, a pose of higher quality than the confidence-selected pick was present within the same 100-pose ensemble. This means that the diffusion-based sampling machinery of both tools can access near-native binding modes for the vast majority of cyclic peptide–protein complexes; the bottleneck lies in identifying which of the sampled modes is correct. This finding resonates with the long-standing recognition in classical molecular docking that decoy generation and scoring are distinct challenges, and that the latter typically limits practical accuracy, and it has recently been recapitulated for AlphaFold-family predictions of protein–protein and linear peptide–protein complexes (Genz et al., 2025; Peng et al., 2025). For cyclic peptides specifically, the problem may be amplified by the small, feature-poor interfaces: ipTM-based scores, which summarize interface quality through a single global similarity metric, are inherently less discriminating between alternative binding registers when the interface comprises only a handful of contact residues. A caveat on what constitutes an incorrect pose is warranted. Proteins and their complexes populate conformational ensembles, and an X-ray structure represents the dominant, crystal-stabilized member of that ensemble; a predicted pose that disagrees with the deposited coordinates is therefore not necessarily wrong: it may correspond to an alternative binding mode that is genuinely populated in solution. Among high-scoring poses (DockQ above 0.80) we cannot distinguish which reproduce the experimentally resolved state and which represent alternative, possibly real, conformations; several may be simultaneously valid. Low-scoring poses, by contrast, are wrong in an unambiguous structural sense: poses below the Acceptable threshold (DockQ below 0.23) place the binder at a median interface RMSD of 8.6–9.6 Å from the native interface (10th–90th percentile 4.0–14.3 Å; Figure S4), a wholesale change of binding register rather than an alternative conformation of the correct one. The Good/Bad labeling used by our rescoring models is thus conservative under the ensemble interpretation: the poses it targets for rejection are structurally displaced from the native state, not merely different from it.

Our feature-decorrelation analysis and machine-learning (ML) experiments demonstrate that the scoring bottleneck can be partially relieved by augmenting native confidence with complementary structural descriptors. The useful descriptors are those carrying incremental signal beyond the confidence axis, not simply those most decoupled from it: per-residue hydrogen-bond density, though moderately correlated with the self-score (|ρ| = 0.32–0.51), was selected for both tools, alongside the essentially orthogonal target electrostatics (Protenix) and the moderately anti-correlated relative interface burial (Boltz), whereas features that cluster tightly with ipTM (pTM, complex pLDDT, interface pLDDT; |ρ| of at least 0.72) contributed little incremental signal once the self-score was included; both selected models improved out-of-fold ROC-AUC (significant for Protenix, p < 0.001; directional but not individually significant for Boltz, p = 0.28). This reliance on physicochemical descriptors external to the predictor’s own score is the same principle underlying knowledge-based interface-affinity predictors such as PRODIGY, which deliberately rely on contact counts and surface-area properties that are independent of any particular docking algorithm’s internal score (Xue et al., 2016). The convergence of both tools’ selected models on per-residue hydrogen-bond density as the single most informative complementary descriptor (Kuroda and Gray, 2016; Pace et al., 2014) points to a systematic blind spot in ipTM-based confidence: such scores register that an interface has formed but do not adequately weight whether the specific polar contacts that define a correct binding register are satisfied. We emphasize that these results are correlative (our ML models establish that the descriptors carry pose-discriminating signal, not that modifying the hydrogen-bond term in the underlying confidence function would necessarily improve prediction), but they identify clear, physically motivated directions for confidence recalibration.

The practical implications for cyclic-peptide structure-based design are twofold. First, researchers using Boltz or Protenix should not rely exclusively on the single top-confidence pose: our data show that a higher-quality conformer is almost always present in the sampled ensemble and can be surfaced by computing a small set of chain-length-normalized interface descriptors (hydrogen-bond density, relative SASA burial, and, for Protenix, target net charge) and ranking poses by a composite score. This rescoring protocol requires no GPU time and only minutes of commodity CPU time: in our runs on a single NVIDIA RTX PRO 5000 Black-well GPU, generating a 100-pose ensemble took about 15 GPU-minutes per target for Boltz-2 and about 2.5 GPU-minutes for Protenix, while computing the full PyMOL and Rosetta descriptor set for the same ensemble took about 10 CPU-minutes (about 1 and about 5 CPU-seconds per pose, respectively) on an Intel Xeon w7-2495X CPU (24 cores; comparable to the prediction step itself rather than negligible, yet trivially parallelizable across poses and targets) and can substantially reduce the risk of carrying a confidently incorrect binding mode into downstream design or optimization (Ji et al., 2024; Rettie et al., 2025). Second, the chemistry- and size-dependent failure modes we identified can inform triage in virtual screening campaigns: disulfide-cyclized and small-target complexes warrant heightened scrutiny or ensemble-level rescoring, while backbone-cyclized peptides on large, well-characterized receptors can be used with greater confidence in the native score. We note, however, that the performance of our feature-based rescoring was established retrospectively, and its behavior in fully prospective design pipelines (where the native structure is unavailable) requires separate validation.

Several limitations should be acknowledged. First, the benchmark comprises 111 targets, which is substantial for cyclic peptide–protein complexes but modest in comparison to linear-peptide or protein–protein interaction benchmarks (Zhai et al., 2025; Peng et al., 2025); the five cyclization classes are imbalanced (SS*1 = 61, SS*3 = 1), which limits statistical power for rare chemistries and means that the disulfide effect is dominated by single-bridge peptides. Second, all pose-quality assessment is funneled through DockQ v2 in CAPRI-peptide mode (Basu and Wallner, 2016; Mirabello and Wallner, 2024), a single composite metric that reduces interface accuracy to a scalar and may not fully capture chemically important features such as side-chain rotamer recovery, water-mediated contacts, or the geometry of non-canonical residues. Third, this is a retrospective study: all 111 targets are deposited in the PDB and may overlap, directly or through homologs, with the training data of both models, potentially inflating absolute accuracy estimates relative to truly novel targets. Prospective validation on newly determined structures, and out-of-distribution evaluation on chemistries absent from this benchmark (e.g., lactam bridges, hydrocarbon staples, N-methylated backbones), will be essential to establish the generalizability of both the observed accuracy trends and the ML-based rescoring strategy. Finally, the rescoring models were trained and evaluated within a single benchmark; their transferability across target classes and their utility as integrated post-processing modules within prediction pipelines remain open questions.

## Methods

### Dataset curation and standardization

The benchmark integrates previously published cyclic peptide–protein complex data (Zhao et al., 2024, 2025) with publicly available Protein Data Bank structures, and comprises 111 non-redundant cyclic peptide–protein complexes. All structures were obtained from the Protein Data Bank (wwPDB consortium, 2019) and standardized as follows. Crystallographic alternate locations were resolved by retaining only the primary conformer (blank or “A”). The benchmark comprises 107 complexes solved by X-ray diffraction, 3 by electron microscopy, and 1 by solution NMR; for the NMR ensemble (7MLA) the first deposited model was taken as the reference conformer. To remove redundant biological or crystallographic copies and leave a single binder–receptor complex per target, multi-copy entries were reduced to the single receptor complex contacted by the binder (six complexes reduced, 105 unchanged). A receptor chain was defined as contacted when any backbone Cα–Cα distance between the binder and that chain fell below 8.0 Å; each standardized complex retains only the receptor chains contacting the binder, and target length was defined as the total number of residues of these receptor chains. All benchmark structures are composed exclusively of natural amino acids; entries containing non-canonical residues were excluded during curation.

Cyclic peptides were classified by chemical connectivity, using the experimental SSBOND records for disulfide bridges and the presence of a head-to-tail N(1)–C(last) amide bond for backbone cyclization. Peptides carrying only the head-to-tail amide bond were classed as backbone-only (BB); those additionally forming one or more intra-peptide disulfide bridges as BB+SS; and those cyclized solely by one, two, or three disulfide bridges as SS*1, SS*2, and SS*3, respectively (Figure 1a). For binary stratification, targets were grouped as “backbone-only” (BB, n = 27) versus “disulfide-cyclized” (BB+SS and SS*1–SS*3 combined, n = 84).

### Structure prediction

Predictions were generated with two deep-learning co-folding methods, Boltz-2 (`boltz2`) and Protenix (base model, `protenix_base_20250630_v1.0.0`). These two configurations were selected from a pilot comparison of seven model–MSA variants (1 seed × 5 samples per configuration; Figures S1 and S2): within each tool family the MSA-enabled variant gave the highest median CAPRI-peptide DockQ, and Protenix collapsed to near-zero DockQ without an MSA whereas Boltz was largely MSA-insensitive, motivating MSA-enabled runs throughout. The pilot confidence–DockQ calibration (Spearman ρ of 0.55 for Boltz and 0.64 for Protenix base) was used to characterize rather than to select the configurations, as model selection prioritized DockQ performance. For each target, 10 random seeds were sampled with 10 conformers per seed, yielding 100 predicted poses per target (11,100 poses per tool; 22,200 poses in total). The receptor chains were supplied with precomputed multiple-sequence alignments generated with MMseqs2 (Steinegger and Söding, 2017) against UniRef90 (Suzek et al., 2015) using the GPU-accelerated AlphaFast parameter profile, which affords about 70-fold faster alignment construction at indistinguishable structural accuracy relative to the default CPU-bound pipeline (Perry et al., 2026); the cyclic-peptide binder, too short to yield a meaningful alignment, was provided in single-sequence mode for both tools. Cyclization chemistry was encoded in each tool’s native input format (format definitions in the respective source repositories (Wohlwend et al., 2024; Protenix Team et al., 2026)). For Boltz-2, head-to-tail backbone cyclization was specified with a `cyclic: true` flag on the peptide entity, and each disulfide bridge as an explicit SG–SG bond-constraint record derived from the experimental SSBOND connectivity and enforced through the model’s diffusion potentials (`–use_-potentials`); Boltz-2 was run with 10 recycling steps, 200 sampling steps, and 10 diffusion samples per seed, and its composite confidence score (0.8 × pLDDT + 0.2 × ipTM) was taken as the model’s self-confidence. For Protenix, backbone cyclization and every disulfide bridge were likewise specified as explicit `covalent_-bonds` records (an N(1)–C(last) bond for head-to-tail closure and an SG–SG bond per disulfide, both derived from the SSBOND records), so in both tools multi-bridge peptides (e.g. cystine knots) retain their exact cross-link pattern. Protenix was run with its default inference settings and MSA features enabled, and its ranking score (a weighted composite of pTM and ipTM with a steric-clash penalty) was taken as the model’s self-confidence. Pose generation averaged about 9 s/pose (Boltz-2) and about 1.5 s/pose (Protenix) on a single NVIDIA RTX PRO 5000 Blackwell GPU (about 15 and about 2.5 GPU-minutes per 100-pose target, respectively); interface-descriptor computation (see below) averaged about 6 CPU-s/pose (PyMOL about 1 s, Rosetta about 5 s) on an Intel Xeon w7-2495X CPU (24 cores/48 threads). Timings are wall-clock estimates derived from run logs and output-file timestamps.

### Pose-quality evaluation

Each predicted pose was scored against its native structure with DockQ v2 in CAPRI-peptide mode, which uses a Cβ-based contact definition adapted to peptide–protein interfaces. Poses were assigned to the standard DockQ tiers: High (0.80 or higher), Medium (0.49 or higher), Acceptable (0.23 or higher), and Incorrect (below 0.23). For binary classification, a pose was defined as Good when its CAPRI-peptide DockQ exceeded 0.80, and Bad otherwise.

### Feature extraction and chain-length normalization

For every pose we assembled a feature vector combining model-internal and externally computed descriptors. Model-internal signals were read directly from each tool’s prediction sidecars: eight for Boltz-2 (self-confidence, pTM, ipTM, complex pLDDT, protein ipTM, complex ipLDDT, predicted distance error, and interface predicted distance error) and five for Protenix (self-confidence, pTM, ipTM, pLDDT, and global predicted distance error). Interface descriptors were computed with PyMOL (Schrödinger LLC, 2025) and Rosetta (Leaver-Fay et al., 2011), following protocols adapted from the ProtDBench evaluation pipeline (Liu et al., 2026), and included solvent-accessible surface area (SASA), interface buried SASA (ΔSASA), the non-polar interface fraction, inter-chain hydrogen bonds, binder-side and target-side interface secondary-structure fractions (assigned with PyMOL’s built-in dss command; shown as “paratope”/”epitope” labels in the figures), the fraction of the binder at the interface, and the sequence net charge and hydropathy of each chain.

Because several of these descriptors scale with complex size, seven size-dependent counts and surface areas were converted to chain-length-normalized densities by dividing by binder length or target length (per-residue SASA and buried SASA; per-residue hydrogen bonds for both PyMOL and Rosetta; and the percentage of the target in the interface). Twelve scale-invariant features were retained unchanged. The remaining raw, size-defining quantities (interface residue counts, non-polar contact counts, and raw hydrogen-bond, SASA, and buried-SASA values from both PyMOL and Rosetta) were represented only through the scale-invariant fractions and per-residue densities described above and were not included as separate features. Together with the model-internal signals this yielded curated pools of 27 features for Boltz-2 and 24 for Protenix.

### Pose-quality classification

A gradient-boosted decision-tree classifier (LightGBM (Ke et al., 2017); 200 estimators, learning rate 0.05, 31 leaves, subsampling rate 0.8, L2 regularization 1.0) was trained to distinguish Good from Bad poses. To prevent leakage across structurally related conformers, evaluation used stratified 5-fold cross-validation with whole targets (PDB IDs) held out together (StratifiedGroupKFold, k = 5, grouped by target). Feature selection used Sequential Forward Floating Selection (Pudil et al., 1994) within an inner grouped 5-fold loop; each tool’s own self-confidence was force-seeded into every model to quantify the incremental contribution of the remaining descriptors, and features selected in at least half of the outer folds formed the consensus model. Model interpretation used TreeSHAP (Lundberg et al., 2020) applied to the consensus features. Confidence intervals and empirical *p*-values for the ROC-AUC gain over the self-confidence baseline were obtained from a 1,000-resample block bootstrap that resampled whole targets. The reported consensus model was then refit on all poses for TreeSHAP interpretation.

### Statistical analysis

Pairwise feature associations were summarized as Spearman rank-correlation coefficients (ρ) pooled across all 11,100 poses per tool; correlation matrices were ordered by average-linkage hierarchical clustering on the distance 1 - |ρ|. Differences in DockQ distributions between target-attribute classes (binder length 10 or fewer vs. more than 10 residues; target length 200 or fewer vs. more than 200 residues; backbone-only vs. disulfide-cyclized) were tested with two-sided Mann–Whitney U tests. Whether the best pose within each tool’s 100-pose pool exceeded its top-confidence pick was assessed with one-sided exact sign tests on the per-target quality gaps.

## Data and code availability

The curated benchmark structures (PDBs), multiple-sequence alignments (.a3m), and all code for dataset preparation, Boltz and Protenix prediction runs, DockQ evaluation, interface-descriptor computation, and rescoring-model building are available at https://github.com/IVB-Generative-Biology/cyclic-benchmark.

## ACKNOWLEDGEMENTS

This work was supported by Pharmaron’s internal R&D grant (Bio-PHAR-YFG-2603002). We thank our colleagues in the Generative Biology Group, In Vitro Biology, Pharmaron, for their input and advice. Generative AI (GLM 5.3) was used to facilitate coding and literature search for this project. AI-generated code and AI-identified literature were manually verified by the authors.

## AUTHOR CONTRIBUTIONS

F.H., P.P. and K.H. secured funding for the project. Y.Y. conceptualized the project. Z.L. conducted benchmarking and modeling with input from Y.Y. and K.H. All authors contributed to manuscript writing.

## Supplementary Information

**Figure S1.**
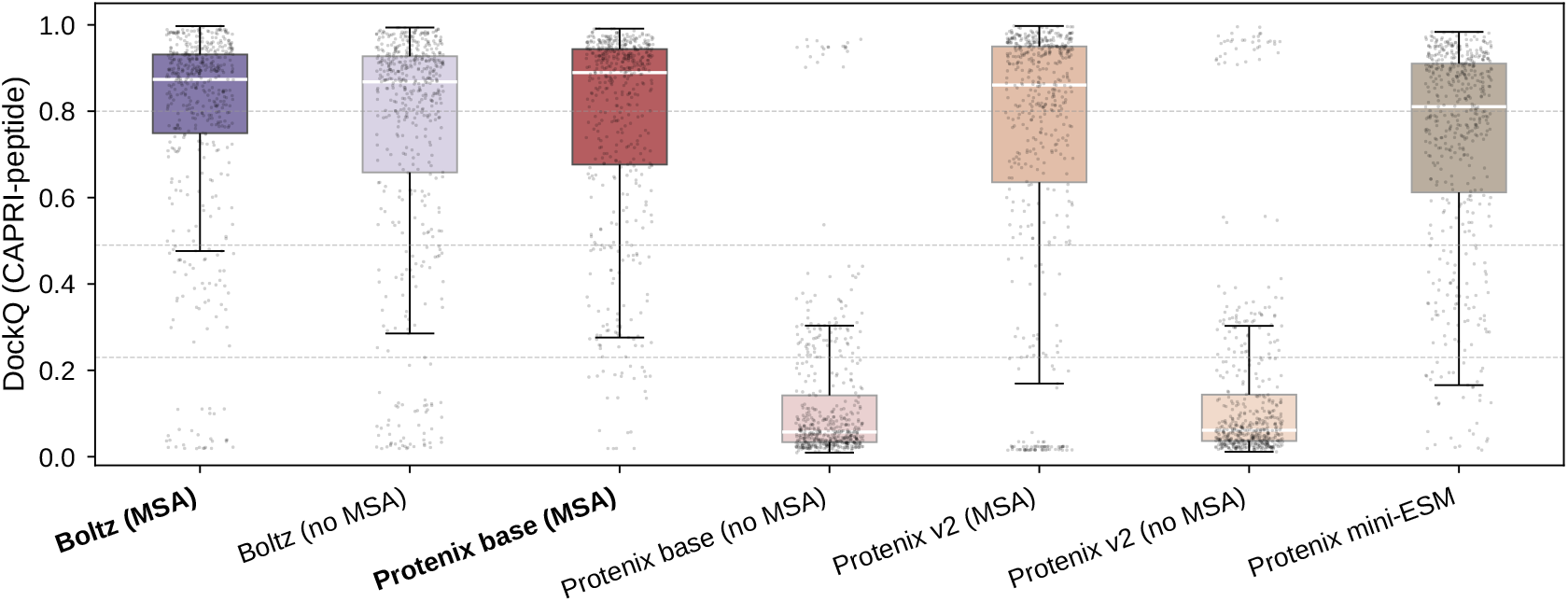
Pilot study: prediction quality across model configurations. Per-pose CAPRI-peptide DockQ distributions for the seven pilot model configurations evaluated while selecting the prediction setup for the main benchmark (1 seed × 5 samples per configuration). The seven configurations are Boltz-2 with and without MSA input; the Protenix base model (`protenix_base_20250630_v1.0.0`; 368 M parameters) with and without MSA; Protenix-v2 (`protenix-v2`; 464 M parameters), the scaled-up Protenix successor with expanded representation dimensionality and further training improvements, with and without MSA; and Protenix-Mini-ESM (`protenix_mini_esm_v0.5.0`; 135 M parameters), a lightweight variant that substitutes ESM2 protein-language-model embeddings for MSA input and was run without MSA. Configurations are grouped by tool (Boltz, purple family; Protenix, red/orange family), and within each tool the MSA-enabled variant is drawn at full saturation and the no-MSA variant lighter. The two configurations adopted for the main benchmark, Boltz with MSA and Protenix base with MSA (bold tick labels, full color), are the top performers within their respective tool families (median DockQ 0.874 and 0.889). Dashed lines mark the DockQ High (0.80), Medium (0.49), and Acceptable (0.23) tiers. Protenix collapses without an MSA input (the base and v2 no-MSA variants fall to near-zero DockQ), whereas the ESM-embedding Mini variant remains serviceable without MSA and Boltz is comparatively MSA-insensitive, motivating the use of MSA-enabled configurations throughout this study.

**Figure S2.**
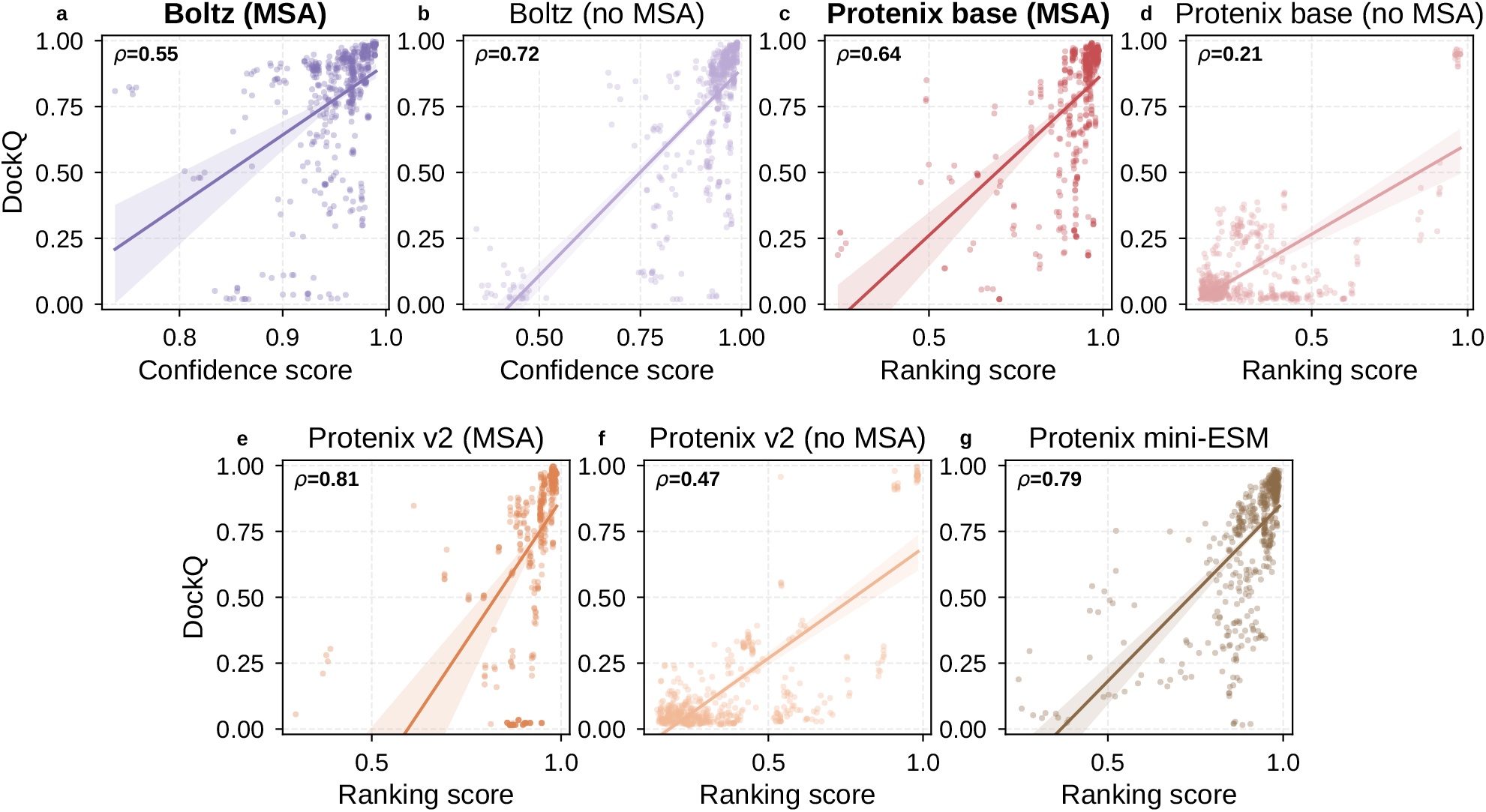
Pilot study: confidence-score calibration across model configurations. Per-pose scatter of CAPRI-peptide DockQ against each configuration’s native self-confidence score (Boltz confidence score, purple; Protenix ranking score, red/orange), with a linear fit; each panel reports the Spearman rank correlation (ρ) (1 seed × 5 samples per configuration). Panels (**a–g**) correspond to the seven pilot configurations of Figure S1, in the same order and colors. The two configurations adopted for the main benchmark, Boltz with MSA (ρ = 0.55) and Protenix base with MSA (ρ = 0.64), are shown in bold. Although Protenix-v2 with MSA (ρ = 0.81) and Mini-ESM (ρ = 0.79) calibrate better than the adopted Protenix base (ρ = 0.64), base with MSA was retained because configuration selection prioritized pose quality (the primary deliverable of prediction) over score calibration, and it was the top performer overall (median CAPRI-peptide DockQ 0.889 vs 0.861 and 0.811; Figure S1); imperfect confidence calibration is precisely the property this study quantifies and mitigates with external descriptors.

**Figure S3.**
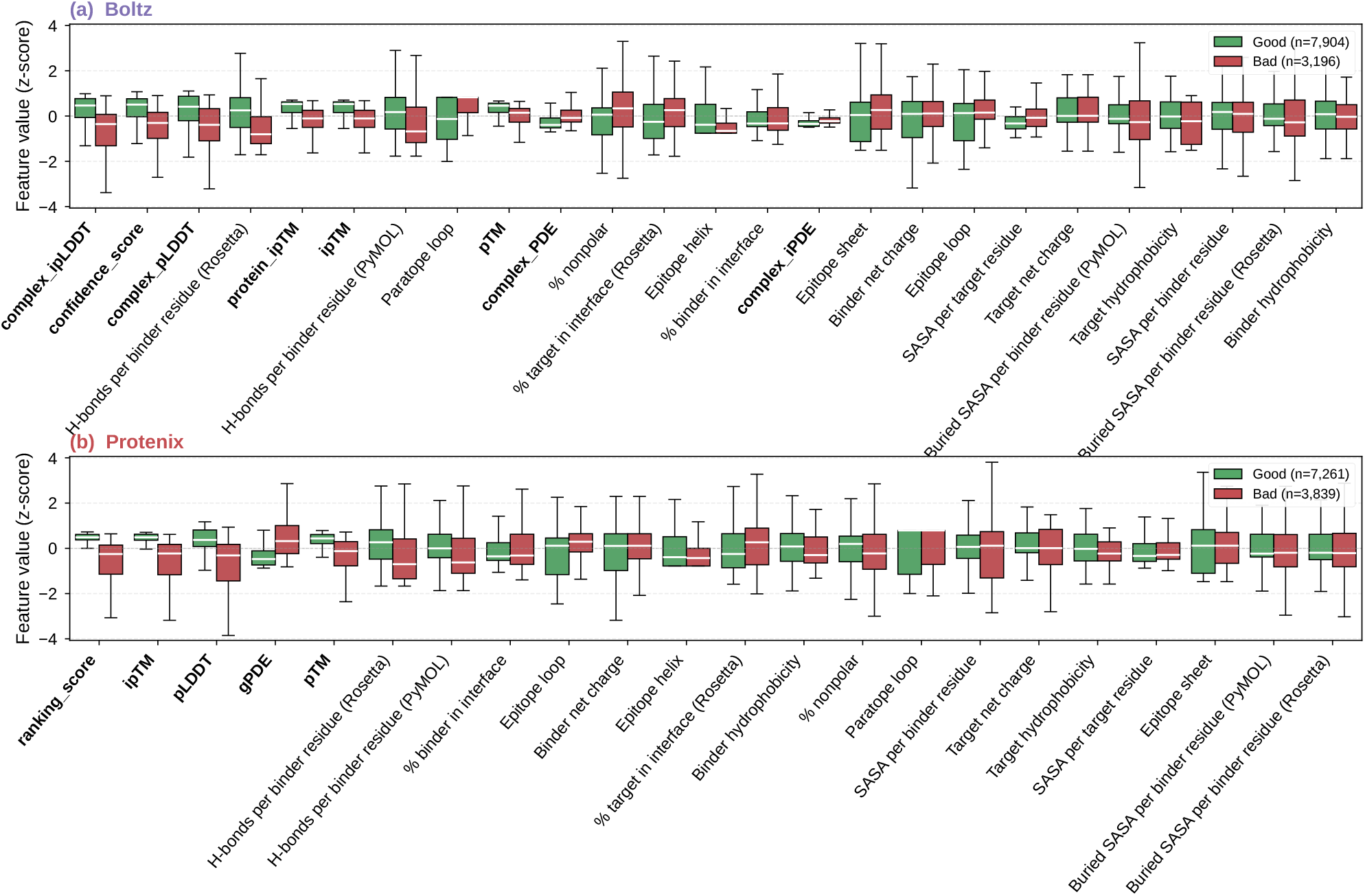
Features separating Good from Bad poses. Per-feature, z-scored boxplots contrasting Good poses (CAPRI-peptide DockQ above 0.80) against Bad poses (DockQ 0.80 or lower), pooled across all 22,200 poses (11,100 per tool; in-panel legend gives each tool’s Good/Bad counts). **(a)** Boltz. **(b)** Protenix. Each feature is standardized to a z-score (pooled over Good+Bad within the tool) so that all descriptors share one y-axis; the shared scale makes the Good/Bad shift directly comparable across features. X-axis labels are bold for model-internal signals and normal for interface descriptors.

**Figure S4.**
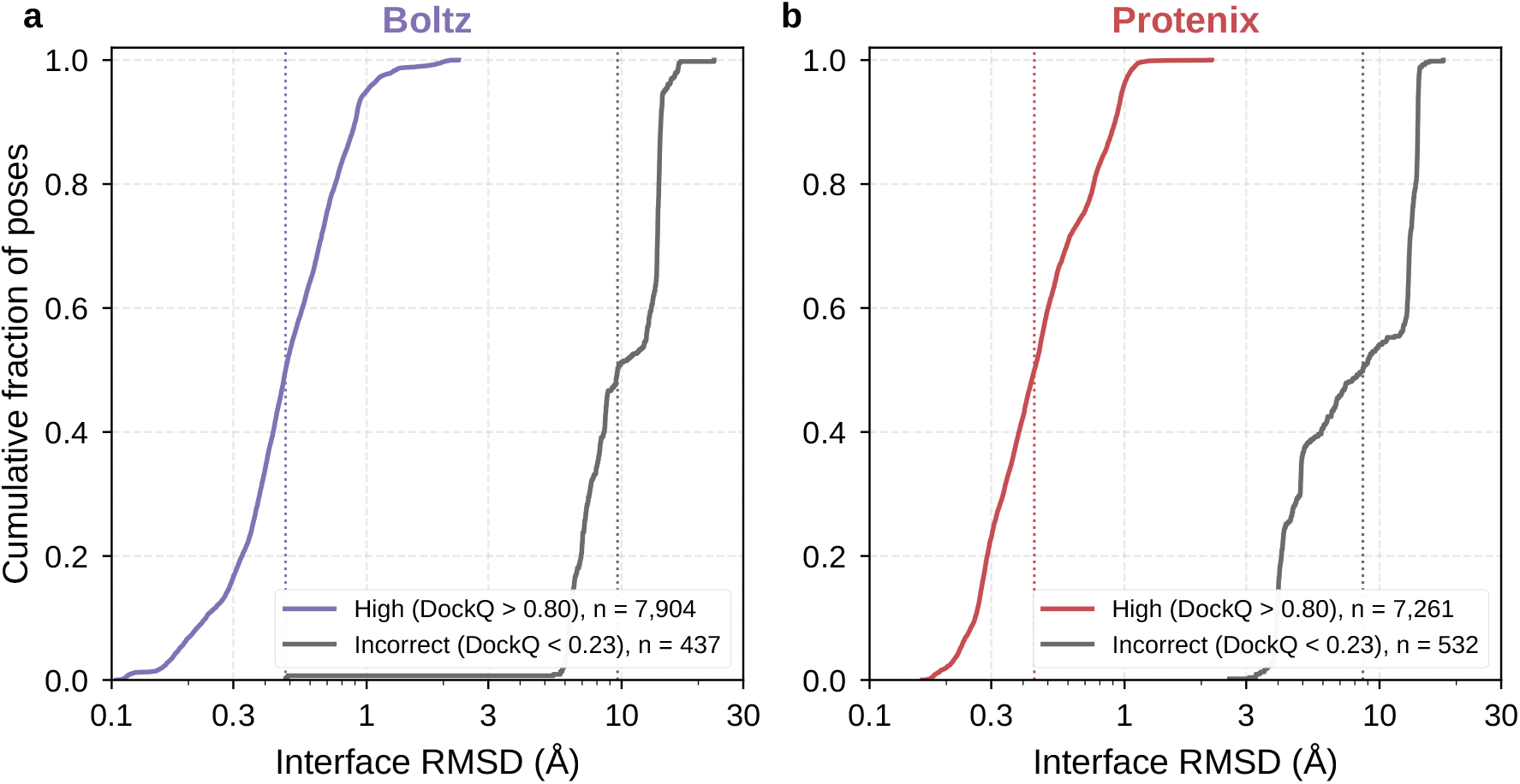
Structural separation between DockQ tiers. Empirical cumulative distributions of interface RMSD (iRMSD, standard DockQ definition) for poses in the High tier (CAPRI-peptide DockQ above 0.80; tool color) versus the Incorrect tier (DockQ below 0.23; gray), pooled over all poses per tool (Boltz, n = 7,904 High / 437 Incorrect; Protenix, n = 7,261 High / 532 Incorrect; note the logarithmic x-axis; dotted vertical lines mark tier medians). High-tier poses cluster at sub-angstrom interface RMSD (median 0.48 Å for Boltz and 0.44 Å for Protenix; 10th–90th percentile 0.23–0.90 and 0.26–0.92 Å), indicating recovery of the native binding register. Incorrect-tier poses sit at median 9.64 Å (Boltz; 10th–90th percentile 6.29–14.26) and 8.61 Å (Protenix; 3.95–14.13), an approximately 20-fold separation consistent with a wholesale change of binding register rather than an alternative conformation of the correct one.

## References

Abramson, J., Adler, J., Dunger, J., Evans, R., Green, T., et al. (2024). Accurate structure prediction of biomolecular interactions with AlphaFold 3. Nature, 630:493–500. doi: 10.1038/s41586-024-07487-w.

Basu, S. and Wallner, B. (2016). DockQ: a quality measure for protein–protein docking models. PLoS ONE, 11:e0161879. doi: 10.1371/journal.pone.0161879.

Chakravarty, D., Guharoy, M., Robert, C. H., Chakrabarti, P., and Janin, J. (2013). Reassessing buried surface areas in protein–protein complexes. Protein Sci., 22:1453–1457. doi: 10.1002/pro.2330.

Evans, R., O’Neill, M., Pritzel, A., Antropova, N., Senior, A., et al. (2021). Protein complex prediction with AlphaFold-Multimer. bioRxiv. doi: 10.1101/2021.10.04.463034.

Genz, L. R., Nair, S., Nagar, N., and Topf, M. (2025). Assessing scoring metrics for AlphaFold2 and AlphaFold3 protein complex predictions. Protein Sci., 34:e70327. doi: 10.1002/pro.70327.

Gerlach, G. J. and Nicoludis, J. M. (2024). KnotFold: improving peptide structure predictions with simulated coevolution. PRX Life, 2:043018. doi: 10.1103/PRXLife.2.043018.

Ji, X., Nielsen, A. L., and Heinis, C. (2024). Cyclic peptides for drug development. Angew. Chem. Int. Ed., 63:e202308251. doi: 10.1002/anie.202308251.

Ke, G., Meng, Q., Finley, T., Wang, T., Chen, W., Ma, W., Ye, Q., and Liu, T.-Y. (2017). LightGBM: a highly efficient gradient boosting decision tree. Adv. Neural Inf. Process. Syst., 30:3146–3154.

Kuroda, D. and Gray, J. J. (2016). Shape complementarity and hydrogen bond preferences in protein–protein interfaces: implications for antibody modeling and protein–protein docking. Bioinformatics, 32:2451–2456. doi: 10.1093/bioinformatics/btw197.

Leaver-Fay, A., Tyka, M., Lewis, S. M., Lange, O. F., Thompson, J., Jacak, R., et al. (2011). ROSETTA3: an object-oriented software suite for the simulation and design of macro-molecules. Methods Enzymol., 487:545–574. doi: 10.1016/B978-0-12-381270-4.00019-6.

Liu, C., Ren, M., Guan, J., Gong, C., Sun, J., Chen, X., and Xiao, W. (2026). ProtDBench: a unified benchmark of protein binder design and evaluation. arXiv. doi: 10.48550/arXiv.2605.04118.

Liu, Z.-L., Hu, J.-H., Jiang, F., and Wu, Y.-D. (2020). CRiSP: accurate structure prediction of disulfide-rich peptides with cystine-specific sequence alignment and machine learning. Bioinformatics, 36:3385–3392. doi: 10.1093/bioinformatics/btaa193.

Lundberg, S. M., Erion, G., Chen, H., DeGrave, A., Prutkin, J. M., Nair, B., Katz, R., Himmelfarb, J., Bansal, N., and Lee, S.-I. (2020). From local explanations to global understanding with explainable AI for trees. Nat. Mach. Intell., 2:56–67. doi: 10.1038/s42256-019-0138-9.

Mirabello, C. and Wallner, B. (2024). DockQ v2: improved automatic quality measure for protein multimers, nucleic acids, and small molecules. Bioinformatics, 40:btae586. doi: 10.1093/bioinformatics/btae586.

Pace, C. N., Fu, H., Lee Fryar, K., Landua, J., Trevino, S. R., et al. (2014). Contribution of hydrogen bonds to protein stability. Protein Sci., 23:652–661. doi: 10.1002/pro.2449.

Passaro, S. et al. (2025). Boltz-2: towards accurate and efficient binding affinity prediction. bioRxiv. doi: 10.1101/2025.06.14.659707.

Peng, C., Ni, W., Liu, Q., Hu, G., and Zheng, W. (2025). A comprehensive benchmarking of the AlphaFold3 for predicting biomacromolecules and their interactions. Brief. Bioinformatics, 26:bbaf616. doi: 10.1093/bib/bbaf616.

Perry, B. C., Kim, J., and Romero, P. A. (2026). AlphaFast: high-throughput AlphaFold 3 via GPU-accelerated MSA construction. bioRxiv. doi: 10.64898/2026.02.17.706409.

Protenix Team, Zhang, Y., Gong, C., Zhang, H., Ma, W., et al. (2026). Protenix-v1: toward high-accuracy open-source biomolecular structure prediction. bioRxiv. doi: 10.64898/2026.02.05.703733.

Pudil, P., Novovičová, J., and Kittler, J. (1994). Floating search methods in feature selection. Pattern Recognit. Lett., 15(11):1119–1125. doi: 10.1016/0167-8655(94)90127-9.

Rettie, S. A., Campbell, K. V., Bera, A. K., Kang, A., et al. (2025). Cyclic peptide structure prediction and design using AlphaFold2. Nat. Commun., 16:4730. doi: 10.1038/s41467-025-59940-7.

Schrödinger LLC. The PyMOL molecular graphics system, (2025). Version 3.1.0.

Steinegger, M. and Söding, J. (2017). MMseqs2 enables sensitive protein sequence searching for the analysis of massive data sets. Nat. Biotechnol., 35(11):1026–1028. doi: 10.1038/nbt.3988.

Suzek, B. E., Wang, Y., Huang, H., McGarvey, P. B., Wu, C. H., and UniProt Consortium. (2015). UniRef clusters: a comprehensive and scalable alternative for improving sequence similarity searches. Bioinformatics, 31(6):926–932. doi: 10.1093/bioinformatics/btu739.

Wohlwend, J., Corso, G., Passaro, S., et al. (2024). Boltz-1: democratizing biomolecular interaction modeling. bioRxiv. doi: 10.1101/2024.11.19.624167.

wwPDB consortium. (2019). Protein Data Bank: the single global archive for 3D macromolecular structure data. Nucleic Acids Res., 47:D520–D528. doi: 10.1093/nar/gky949.

Xie, X., Li, C. Z., Lee, J. S., and Kim, P. M. (2025). CyclicBoltz1, fast and accurately predicting structures of cyclic peptides and complexes containing non-canonical amino acids using AlphaFold 3 framework. bioRxiv. doi: 10.1101/2025.02.11.637752.

Xue, L. C., Rodrigues, J. P. G. L. M., Kastritis, P. L., Bonvin, A. M. J. J., and Vangone, A. (2016). PRODIGY: a web server for predicting the binding affinity of protein–protein complexes. Bioinformatics, 32:3676–3678. doi: 10.1093/bioinformatics/btw514.

Zhai, S., Zhao, H., Wang, J., Lin, S., Liu, T., Jiang, D., Liu, H., Kang, Y., Yao, X., and Hou, T. (2025). PepPCBench is a comprehensive benchmark for protein-peptide complex structure prediction with AlphaFold3. bioRxiv. doi: 10.1101/2025.04.08.647699.

Zhang, C., Wang, W., Zhu, N., et al. (2025). AlphaFold3 for noncanonical cyclic peptide modeling: hierarchical benchmarking reveals accuracy and practical guidelines. J. Chem. Inf. Model., 65:9777–9789. doi: 10.1021/acs.jcim.5c01393.

Zhang, H. and Chen, S. (2022). Cyclic peptide drugs approved in the last two decades (2001–2021). RSC Chem. Biol., 3:18–31. doi: 10.1039/d1cb00154j.

Zhao, H., Jiang, D., Shen, C., et al. (2024). Comprehensive evaluation of 10 docking programs on a diverse set of protein–cyclic peptide complexes. J. Chem. Inf. Model., 64: 2112–2124. doi: 10.1021/acs.jcim.3c01921.

Zhao, H., Huang, J., Weng, G., Jiang, D., Hu, R., Kang, Y., and Hou, T. (2025). Improving the predictive performance of binding affinities and poses for protein–cyclic peptide complexes through fine-tuned MM/PBSA(GBSA)-based methods. Brief. Bioinformatics, 26: bbaf632. doi: 10.1093/bib/bbaf632.

